# An epigenome-wide association study of educational attainment (*n* = 10,767)

**DOI:** 10.1101/114637

**Authors:** Richard Karlsson Linnér, Riccardo E Marioni, Cornelius A Rietveld, Andrew Simpkin, Neil M Davies, Kyoko Watanabe, Nicola J Armstrong, Kirsi Auro, Clemens Baumbach, Marc Jan Bonder, Jadwiga Buchwald, Giovanni Fiorito, Khadeeja Ismail, Stella Iurato, Anni Joensuu, Pauliina Karell, Silva Kasela, Jari Lahti, Allan F McRae, Pooja R Mandaviya, Ilkka Seppälä, Yunzhang Wang, Laura Baglietto, Elisabeth B Binder, Sarah E Harris, Allison M Hodge, Steve Horvath, Mikko Hurme, Magnus Johannesson, Antti Latvala, Karen A Mather, Sarah E Medland, Andres Metspalu, Lili Milani, Roger L Milne, Alison Pattie, Nancy L Pedersen, Annette Peters, Silvia Polidoro, Katri Räikkönen, Gianluca Severi, John M Starr, Lisette Stolk, Melanie Waldenberger, BIOS Consortium, Johan G Eriksson, Tõnu Esko, Lude Franke, Christian Gieger, Graham G Giles, Sara Hägg, Pekka Jousilahti, Jaakko Kaprio, Mika Kähönen, Terho Lehtimäki, Nicholas G Martin, Joyce B. C van Meurs, Miina Ollikainen, Markus Perola, Danielle Posthuma, Olli T Raitakari, Perminder S Sachdev, Erdogan Taskesen, André G Uitterlinden, Paolo Vineis, Cisca Wijmenga, Margaret J Wright, Caroline Relton, George Davey Smith, Ian J Deary, Philipp D Koellinger, Daniel J Benjamin

## Abstract

The epigenome has been shown to be influenced by biological factors, such as disease status, and environmental factors, such as smoking, alcohol consumption, and body mass index. Although there is a widespread perception that environmental influences on the epigenome are pervasive and profound, there has been little evidence to date in humans with respect to environmental factors that are biologically distal. Here, we provide evidence on the associations between epigenetic modifications—in our case, CpG methylation—and educational attainment (EA), a biologically distal environmental factor that is arguably among of the most important life-shaping experiences for individuals. Specifically, we report the results of an epigenome-wide association study meta-analysis of EA based on data from 27 cohort studies with a total of 10,767 individuals. While we find that 9 CpG probes are significantly associated with EA, only two remain associated when we restrict the sample to never-smokers. These two are known to be strongly associated with maternal smoking during pregnancy, and thus their association with EA could be due to correlation between EA and maternal smoking. Moreover, their effect sizes on EA are far smaller than the known associations between CpG probes and biologically proximal environmental factors. Two analyses that combine the effects of many probes—polygenic methylation score and epigenetic-clock analyses—both suggest small associations with EA. If our findings regarding EA can be generalized to other biologically distal environmental factors, then they cast doubt on the hypothesis that such factors have large effects on the epigenome.

## Introduction

The epigenome is known to be influenced by biological factors such as disease status^1,2^. There is a widespread perception that the epigenome is also affected by a variety of social environmental factors^3–10^, but virtually all of the replicated evidence to date in humans relates to environmental factors that have a fairly direct biological impact, such as smoking^11–13^, alcohol consumption^14,15^, and excess energy intake resulting in increased body mass index (BMI)^16,17^. Here we study the associations between epigenetic modifications—specifically, the methylation of cytosine-guanine pairs connected by a phosphate link (CpG methylation)—and educational attainment (EA). EA is biologically distal, and yet it is arguably among the most important life-shaping experiences for individuals in modern societies. EA therefore provides a useful test case for whether and to what extent biologically distal environmental factors may affect the epigenome.

In this paper, we report the results of a large-scale epigenome-wide association study (EWAS) meta-analysis of EA. By meta-analysing harmonised EWAS results across 27 cohort studies, we are able to attain an overall sample size of 10,767 individuals, making this study one of the largest EWAS to date^13,15,18^. A large sample size is important because little is known about plausible EWAS effect sizes for complex phenotypes such as EA, and an underpowered analysis would run a high risk of both false negatives and false positives^19,20^.

As is standard in EWAS, we use data on CpG DNA methylation. This is the most widely studied epigenetic mark in large cohort studies^1^. Methylation level is measured by the beta value, which is the proportion of methylated molecules at each CpG site, a continuous variable ranging between 0 and 1^21^. The Illumina 450k Bead Chip measures methylation levels at over 480,000 sites in human DNA and has been used in many cohort studies^1^.

We report results from both of the two common approaches for the analysis of such methylation datasets. Our main analysis is an EWAS, which considers regression models for each CpG methylation site with EA. We also present evidence from an ‘epigenetic clock^,22,23^ analysis, which uses a weighted linear combination of probes (i.e. measured CpG methylation sites) to predict an individual's so-called ‘biological age.’ The resulting variable can then be linked to phenotypes and health outcomes. EWAS studies to date have found associations between DNA methylation and, for example, smoking^11,12^, body mass index (BMI)^16,24^, traumatic stress^25^, alcohol consumption^14,26^, and cancer^2,27^. In prior work, an age-accelerated epigenetic clock (i.e. an older biological than chronological age) has been linked to increased mortality risk^28^, poorer cognitive and physical health^29^, greater Alzheimer’s disease pathology^30^, Down’s syndrome^31^, high lifetime stress^32^, and lower income^33^.

## Methods

### Participating cohorts

Data for the EWAS and epigenetic clock analyses were available in 27 independent cohort studies from 15 cohorts from across Europe, the US, and Australia (**Supplementary Table S1.1**), and the total sample size comprised 10,767 individuals.

### Ethics and preregistration

All contributing data were collected and analysed according to cohort-specific Local Research Ethics Committees or Institutional Review Boards, and the analyses were performed in accordance with a pre-registered analysis plan archived at Open Science Framework (OSF) in September 2015 (available at: https://osf.io/9v3nk/). All participants provided written informed consent.

### Educational attainment measures

Following earlier work of the Social Science Genetics Association Consortium (SSGAC)^34,35^, EA was harmonized across cohorts. The EA variable is defined in accordance with the ISCED 1997 classification (UNESCO), leading to seven categories of EA that are internationally comparable. The categories are translated into US years-of-schooling equivalents, which have a quantitative interpretation (**Supplementary Table S1.2-S1.3**).

### Participant inclusion criteria

To be included in the current analysis, participants had to satisfy six criteria: 1) participants were assessed for educational attainment at or after 25 years of age; 2) participants were of European ancestry; 3) all relevant covariate data were available for each participant; 4) participants passed the cohort-level methylation quality control; 5) participants passed cohort-specific standard quality controls (for example, genetic outliers were excluded); and 6) participants were not disease cases from a case/control study.

### DNA methylation measurement and cohort-level quality control

Whole-blood DNA CpG methylation was measured genome-wide in all cohorts using the Illumina 450k Human Methylation chip. Cohort-specific information regarding technical treatment of the data, such as background-correction^36^, normalisation^37^ and quality control, is reported in **Supplementary Table S1.4.**

### Statistical analysis

#### Epigenome-wide association study (EWAS)

We first ran an EWAS of EA to investigate associations with individual methylation markers (**Supplementary Note 1.4**). As is standard, the EWAS was performed as a set of linear regressions, one methylation marker at a time, with methylation beta value (0-1) as the dependent variable. The key independent variable was EA. We estimated two regression models that differ in the set of covariates included. In the *basic model*, the covariates were age, sex, imputed or measured white blood-cell counts, technical covariates from the methylation array, and four genetic principal components to account for population stratification. In the *adjusted model,* we additionally controlled for body mass index (BMI, kg/m^2^), smoker status (three categories: current, previous, or never smoker), an interaction term between age and sex, and a quadratic term for age. Since BMI and smoking are correlated with EA^38,39^ and known to be associated with methylation^13,17^, the basic model may identify associations with EA that are actually due to BMI or smoking. While the adjusted model reduces that risk, it may also reduce power to identify true associations with EA (by controlling for factors that are correlated with EA). While we present the results for both models, we focus on the adjusted model because it is more conservative. Details of cohort-specific covariates are presented in **Supplementary Table S1.4**.

#### EWAS quality control and meta-analysis

Each participating cohort uploaded EWAS summary statistics to a central secure server for quality control (QC) and meta-analysis. The number of CpG probes filtered at each step of the QC is presented in **Supplementary Table S1.5.** We removed: probes with missing *P*-value, standard error, or coefficient estimate; probes with a call rate less than 95%; probes with a combined sample size less than 1,000; probes not available in the probe-annotation reference by Price et al. (2013)^40^; CpH probes (H = A/C/T); probes on the sex chromosomes; and cross-reactive probes highlighted in a recent paper by Chen et al.^41^. We performed a sample-size weighted meta-analysis of the cleaned results using METAL^42^. We used single genomic control, as is common in genome-wide association studies (GWAS)^43^, to correct the meta-analysis *P*-values for possible unaccounted-for population stratification. Probes with a *P*-value less than 1×10^−7^, a commonly-used threshold in EWAS^1^, were considered epigenome-wide significant associations.

#### Polygenic predictions with polygenic methylation score

Analogous to polygenic-score prediction analyses from GWAS, a polygenic methylation prediction analysis was performed (**Supplementary Note 1.7**). We tested predictive power in three independent cohort studies: Lothian Birth Cohort 1936 (LBC1936, *n* = 918), RS-BIOS (Rotterdam Study - BIOS, *n* = 671), and RS3 (Rotterdam Study 3, *n* = 728). We re-ran the EWAS meta-analysis three times, each time holding out one of the prediction samples. In each prediction sample, we constructed each polygenic methylation score (PGMS) as a weighed sum of the methylation markers’ beta values, using the *Z*-statistics from the EWAS as weights. The *Z*-statistics are used instead of the EWAS coefficients because CpG methylation is the dependent variable in the EWAS regression. We constructed PGMSs using two different thresholds for probe inclusion, *P* < 1 × 10^−5^ and *P* < 1 × 10^−7^, and using the EWAS results from the basic and adjusted EWAS models, for a total of four PGMSs in each prediction sample.

To shed light on the direction of causation of epigenetic associations, we used a fourth prediction cohort study, a sample of children in the ALSPAC ARIES cohort^44^. We constructed a PGMS using the same approach as described above, in this case using data from cord-blood-based DNA methylation at birth. The outcome variables in this sample are average educational achievement test scores (Key Stages 1–4^45^) from age 7 up to age 16 years.

For both our adult and child samples, we measure the predictive power of the PGMS by examining the incremental coefficient of determination (incremental *R*^2^) from adding the PGMS to the regression model for predicting EA (or test scores), i.e., the difference in *R^2^* between the regression model without the PGMS and the regression model with the PGMS added as a predictor.

To enable us to examine the relationship between epigenetic and genetic associations, we also constructed single-nucleotide polymorphism (SNP) polygenic scores for EA. To do so, we used SNP genotype data available in the three adult prediction cohort studies (LBC1936, RS-BIOS, and RS3). We constructed a polygenic score in each sample as a weighted sum of genotypes from all available genotyped SNPs, with GWAS meta-analysis coefficients as weights. We obtained the coefficients by re-running the largest GWAS meta-analysis to date of EA^35^ after excluding the LBC1936, RS-BIOS, and RS3 samples, respectively. We examine the incremental *R^2^* from adding the PGMS to a regression model for predicting EA that already includes the polygenic score. We then re-estimate the model after we add an interaction term between the PGMS and the polygenic score, and we assess the incremental *R^2^* compared to the model with the PGMS and the polygenic score as additive main effects.

#### Epigenetic clock analyses

To construct our epigenetic clock variables, we entered the raw beta-value data into the online Horvath calculator^23^, as per our pre-registered analysis plan. The “normalize data” and “advanced analysis for Blood Data” options were selected. The following variables were selected from the calculator’s output for subsequent analysis:

- Clock 1. Horvath age acceleration residuals, which are the residuals from the regression of chronological age on Horvath age.
- Clock 2. White blood cell count adjusted Horvath age acceleration, which is the residual from Clock 1 after additional covariate adjustment for imputed white blood cell counts.
- Clock 3. White blood cell count adjusted Hannum age acceleration, which is the same as Clock 2 but with the Hannum age prediction in place of the Horvath prediction.
- Clock 4. Cell-count enriched Hannum age acceleration, which is the basic Hannum predictor plus a weighted average of aging-associated cell counts. This index has been found to have the strongest association with mortality^46^.

These Clock measures are annotated in the Horvath software as follows: ‘AgeAccelerationResidual’, ‘AAHOAdjCellCounts’, ‘AAHAAdjCellCounts’, and ‘BioAge4HAAdjAge’. We analysed two regression models, both with EA as the dependent variable and a clock variable as an independent variable. In the *basic age acceleration model,* we control for chronological age, and in the *adjusted age acceleration model,* we additionally control for BMI and smoker status (current, previous, or never smoker). In total, in each adult cohort, we estimated eight regressions: each of the two models with each of the four clock variables as an independent variable. For each of the eight regressions, we performed a sample-size weighted meta-analysis of the cohort-level results.

## Results

### Descriptive statistics

Summary statistics from the 27 independent cohort studies from the 15 contributing cohorts are shown in **Supplementary Table S1.1**. The mean age at reporting ranges from 26.6 to 79.1 years, and the sample size ranges from 48 to 1,658, with a mean of 399 individuals per cohort. The mean cohort EA ranges from 8.6 to 18.3 years of education, and the sample-size weighted mean is 13.6 (SD = 3.62). The meta-analysis sample is 54.1% female.

### EWAS

#### EWAS Quality Control (QC)

The QC filtering is reported in **Supplementary Table S1.5**. We inspected the quantile-quantile (QQ) plot of the filtered EWAS results from each contributing cohort as part of the QC procedure before meta-analysis. The genomic inflation factor (*λ*_*GC*_), defined as the ratio of the median of the empirically observed chi-square test statistics to the expected median under the null, has a mean across the cohorts of 1.02 for the adjusted model (SD = 0.18). We report the cohort-level genomic inflation factor after probe filtering in **Supplementary Table S1.5.** The variation in *λ*_*GC*_ across cohorts is comparable to that from EWAS performed in cohorts of similar sample size^12^. We applied genomic control at the cohort level, which is a conservative method of controlling for residual population stratification that may remain even despite the regression controls for principal components^43^. The meta-analysis *λ*_*GC*_ is 1.19 for the basic model and 1.06 for the adjusted model.

#### EWAS results

**Figure 1** shows the Manhattan plot for the meta-analysis results of the adjusted model. The Manhattan plot for the basic model, as well as the QQ plots for the basic and adjusted model, are in the **Supplementary Note**. In the basic model there are 37 CpG probes associated with EA at our preregistered epigenome-wide *P*-value threshold (*P* < 10^−7^); these results are reported in Supplementary Table S1.6a. In the adjusted model there are 9 associated probes, listed in Table 1 (with additional details in **Supplementary Table S1.7a**), all of which were also associated in the basic model. We hereafter refer to the adjusted model’s 9 associated probes as the “lead probes.”

**Figure 1.**
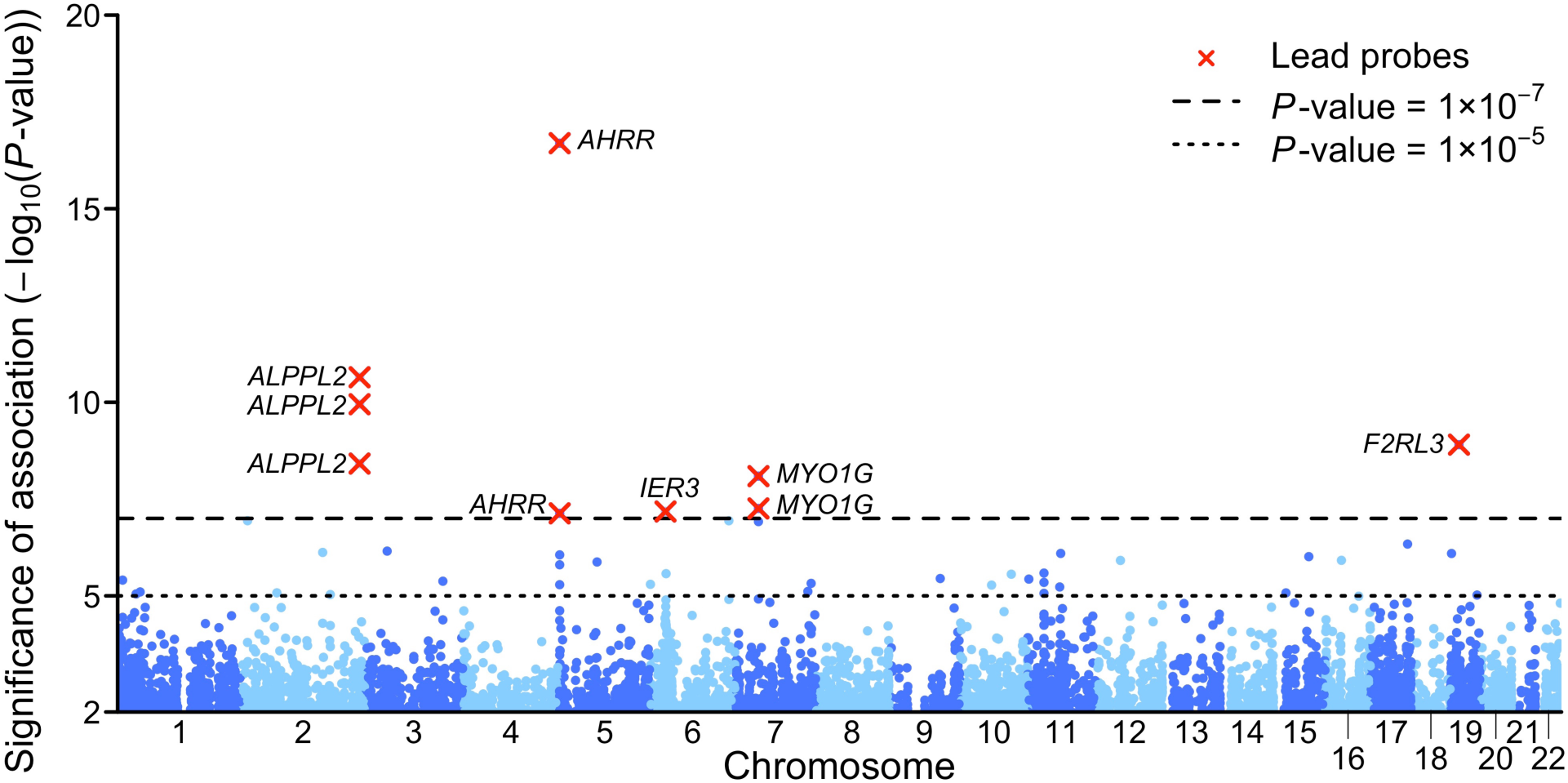
Manhattan plot of the adjusted EWAS model. The figure displays the Manhattan plot of the meta-analysis of the adjusted EWAS model (the Manhattan plot of the basic model is reported in **Supplementary Note**). The *x*-axis is chromosomal position, and the *y*-axis is the significance on a −log_10_ scale. The dashed lines mark the threshold for epigenome-wide significance (*P* = 1×10^−7^) and for suggestive significance (*P* = 1×10^−5^). Each epigenome-wide associated probe is marked with a red ×, and the symbol of the closest gene based on physical position.

**Table 1.**
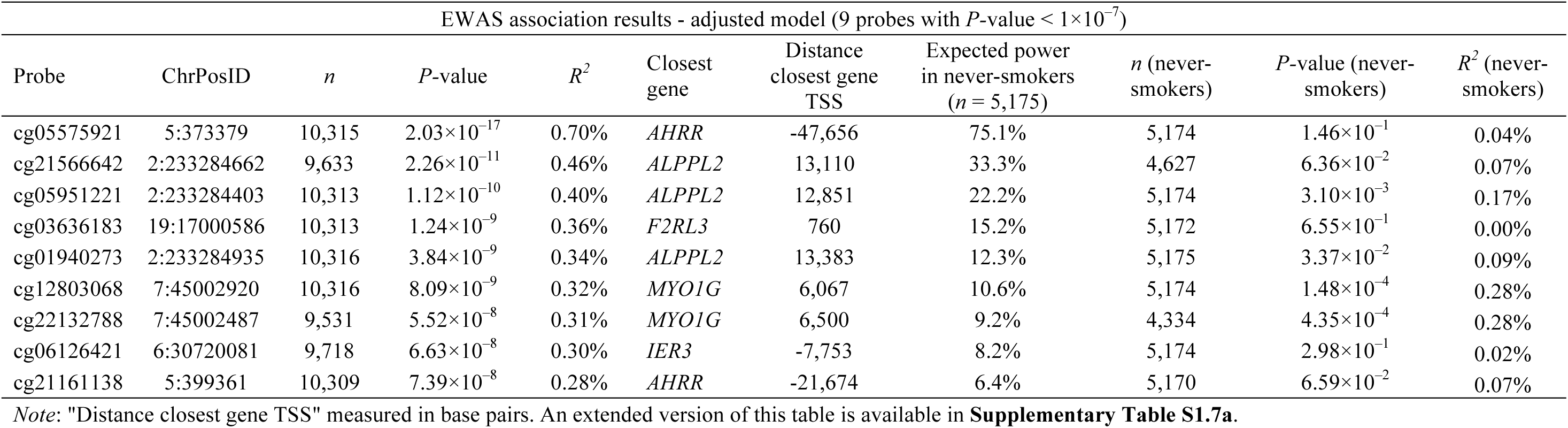
EWAS association results - adjusted model

The effect sizes of the associations for the 9 lead probes are shown in **Table 1**. The coefficients of determination (*R*^2,^s) range from 0.3% to 0.7%. To put these effect sizes in perspective, Figure 2 and Supplementary Table S1.8 compare the *R*^2,^s for the top 50 probes in our adjusted model with the top 50 probes from recent large-scale EWAS of smoking^13^, maternal smoking^12^, alcohol consumption^15^, and BMI^17^, as well as the top 50 GWAS SNP associations with EA^35^. The EA EWAS associations we find are an order of magnitude larger than the largest EA SNP effect sizes from the GWAS. However, our EWAS associations for EA are small in magnitude relative to the EWAS associations that have been reported for more biologically proximal environmental factors. BMI is the most similar to EA, with *R*^2,^s of associated probes approximately 20-50% larger than those for EA. Relative to the largest *R*^2^ for an EA-associated probe, the largest for probes associated with smoking and maternal smoking are greater by factors of roughly 3 and 17, respectively.

**Figure 2.**
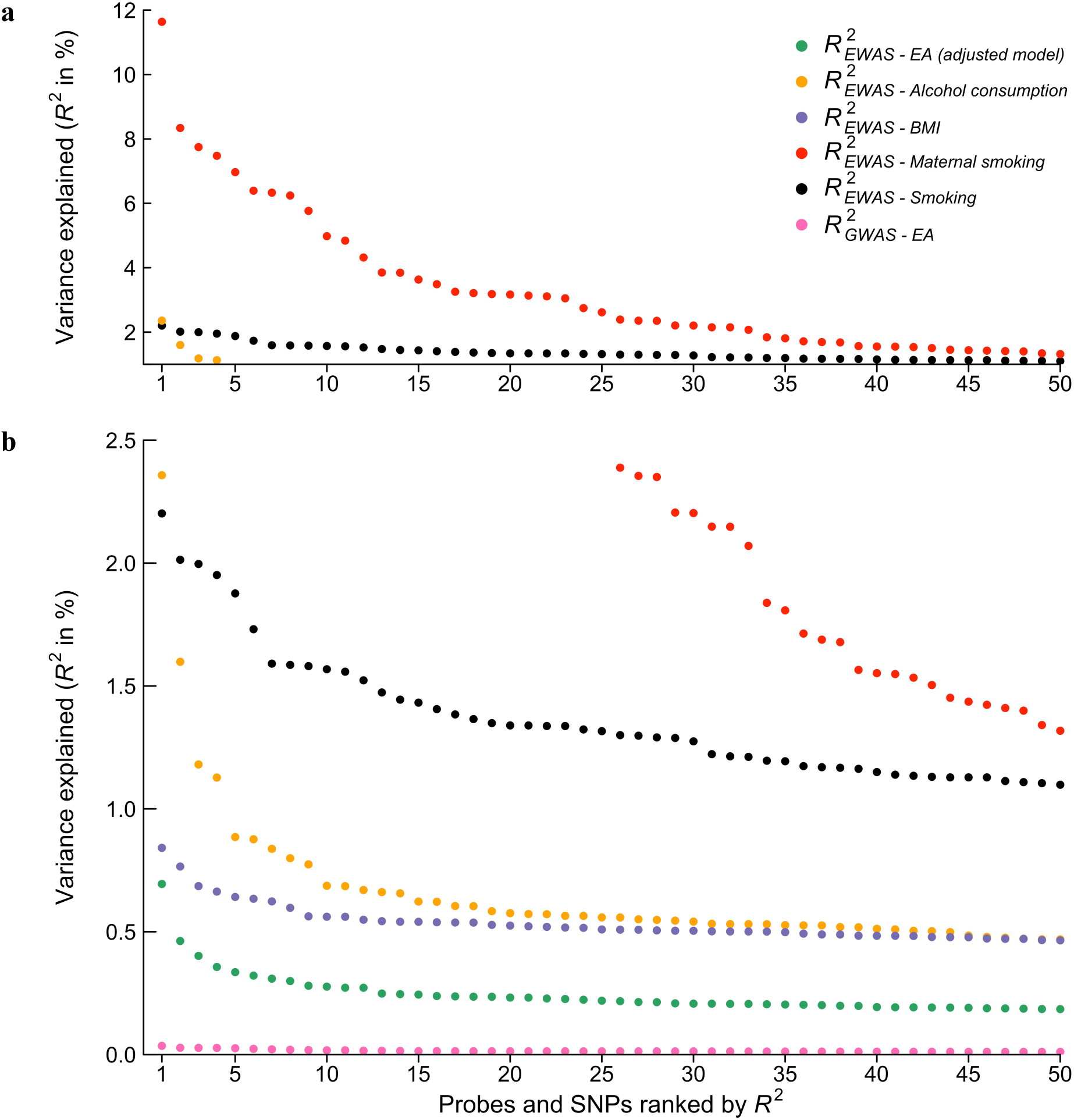
Distribution of coefficient of determination (*R*^2^’s) across traits and methods. The figure displays the effect size estimates in terms of *R^2^*, in descending order, for the 50 top probes of the adjusted EWAS model. For comparison we present the 50 top probes from recent EWAS on alcohol consumption (*n* = 9,643, Liu et al., 2016), BMI (*n* = 7,798, Mendelson et al., 2017), smoking (*n* = 9,389, Joehanes et al., 2016), and maternal smoking (*n* = 6,685, Joubert et al., 2016). For comparison with GWAS effect sizes we contrast the EWAS probes with the effect sizes of the 50 top approximately independent SNPs from a recent GWAS on educational attainment (*n* = 405,073, Okbay et al., 2016). Panel (a) and (b) display the same results but with a different scaling of the *y*-axis in order for the smaller effect sizes to be visible.

#### Lookup of lead probes in published EWAS of smoking

Since our smoker-status control variable is coarse and discrete (current, former, or never smoker), we were concerned that the adjusted EWAS model might not have adequately controlled for exposure to smoking, i.e., the amount and duration of smoking. Therefore, we performed a lookup of our lead probes in published EWAS of smoking (**Supplementary Note 2.3**). The literature review is reported in **Supplementary Table S1.9**. We found that all 9 lead probes have previously been associated with smoking in at least one study. This evidence, however, is difficult to interpret because: (a) since not all studies included EA as a covariate, it is possible that these associations are truly due to a relationship with EA rather than with smoking, and (b) many of the published smoking EWAS are based on small samples and hence likely have an inflated rate of spurious associations that could include EA-associated probes just by chance. Nonetheless, the results of this literature review motivated our analysis of the never-smoker subsample, discussed next.

#### Robustness of EWAS results in the never-smoker subsample

To minimize the possible confounding effect of smoking on the association between EA and CpG methylation, we conducted a set of analyses that we did not anticipate when we preregistered our analysis plan. Specifically, we went back to the cohorts and asked them to re-conduct their EWAS, this time restricting the analysis to individuals that self-reported as never smokers. After following the same QC steps as above, we performed a new meta-analysis of these results (*n* = 5,175).

After re-estimation, none of the 9 lead probes are associated at the stringent epigenome-wide significance threshold (*P* < 10^−7^). However, we strongly reject the joint null hypothesis that none of the 9 probes is associated with EA (one-tailed *P*-value of 2.18×10^−11^; for more details, see **Supplementary Note 2.3.2**). Relative to their effect sizes in the full EWAS sample, in the never-smoker subsample 7 out of the 9 probes have effect sizes (in terms of *R^2^)* that are reduced by more than 60% (see **Table 1** and panel A in **Figure 3**). In contrast, the effect sizes of two probes, cg12803068 and cg22132788, exhibit little or no attenuation. These are the probes for which the evidence of association with EA remains strongest (*P* = 1.48×10^−4^ and *P* = 4.35×10^−4^, respectively).

**Figure 3.**
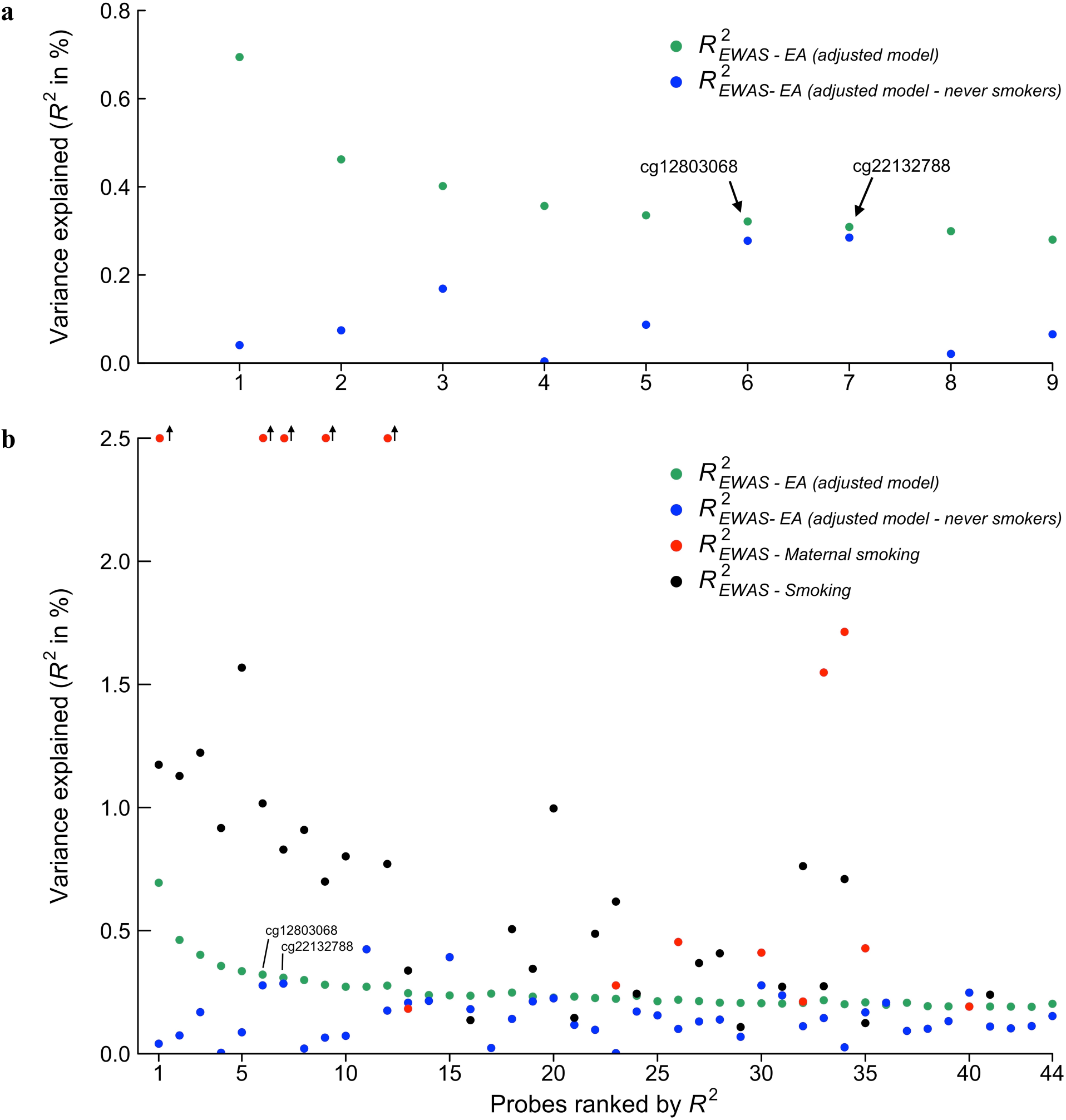
Distribution of *R*^2^ to compare the effect sizes across never smokers and smoking EWAS. Panel **(a)** displays the effect-size estimates in terms of *R*^2^ for the 9 lead probes, in descending order, and the lead probe’s corresponding effect size when re-estimated in the subsample of never smokers. Panel **(b)** displays the same information for the probes of the adjusted model with *P* < 1 × 10"^5^ (including the 9 lead probes), as well as the same probes’ effect-size estimates from two recent EWAS of smoking (*n* = 9,389, Joehanes et al., 2016), and maternal smoking (*n* = 6,685, Joubert et al., 2016). The smoking and maternal smoking estimates are only publicly available for probes associated at FDR < 0.05 in the respective EWAS.

These two probes, however—both in proximity to the gene *MYO1G*—have been found to be associated with maternal smoking during pregnancy, and the effects on the methylation of this gene are persistent when measured at age 17 in the offspring^12,47^. We cannot distinguish between the hypothesis that these probes have some true association with EA and the hypothesis that their apparent association with EA is entirely driven by more maternal smoking during pregnancy among lower-EA individuals. We also cannot rule out that the probes’ association with EA is driven by second-hand smoke exposure, which could also be correlated with EA.

To assess the how widely such confounding may affect the EA results, in Panel B of **Figure 3** we compare the effect sizes of all the probes associated with EA at *P* < 10^−5^ in the adjusted EWAS model to the effect sizes found for the same probes in EWAS meta-analyses of smoking^13^ and maternal smoking^12^ (see also **Supplementary Table S.1.10**). Since effect size estimates for smoking and maternal smoking were not available to us for all the probes, and since all the effect sizes are estimated with error, we cannot draw firm conclusions. Nonetheless, the comparison strongly suggests that residual smoking exposure (i.e., the misclassification of amount and duration of smoking, and second-hand smoke that is not captured by the smoking covariate), and maternal smoking remain potential confounders for the probe associations with EA, even in the subsample of individuals who are self-reported never-smokers.

#### Prediction using polygenic methylation scores

The incremental *R^2^*’s from the prediction of EA with polygenic methylation scores (PGMSs) in our adult prediction cohort studies, the LBC1936, RS-BIOS, and RS3, are reported in **Supplementary Table S1.10a** and in **Figure 4**. Across the four PGMSs constructed from the basic and adjusted model, constructed with estimates from the full EWAS sample, and from both probe-inclusion thresholds (*P* < 10^−5^ and *P* < 10^−7^), the incremental *R*^2^’s range from 1.4% to 2.0% (*P* < 3.28×10^−8^ and lower). There is also weak evidence for an interaction between the PGMS and the polygenic score in predicting EA, with *R^2^*’s for the interaction term ranging from 0.1% to 0.3% (*P*-values ranging from 0.01 to 0.12).

**Figure 4.**
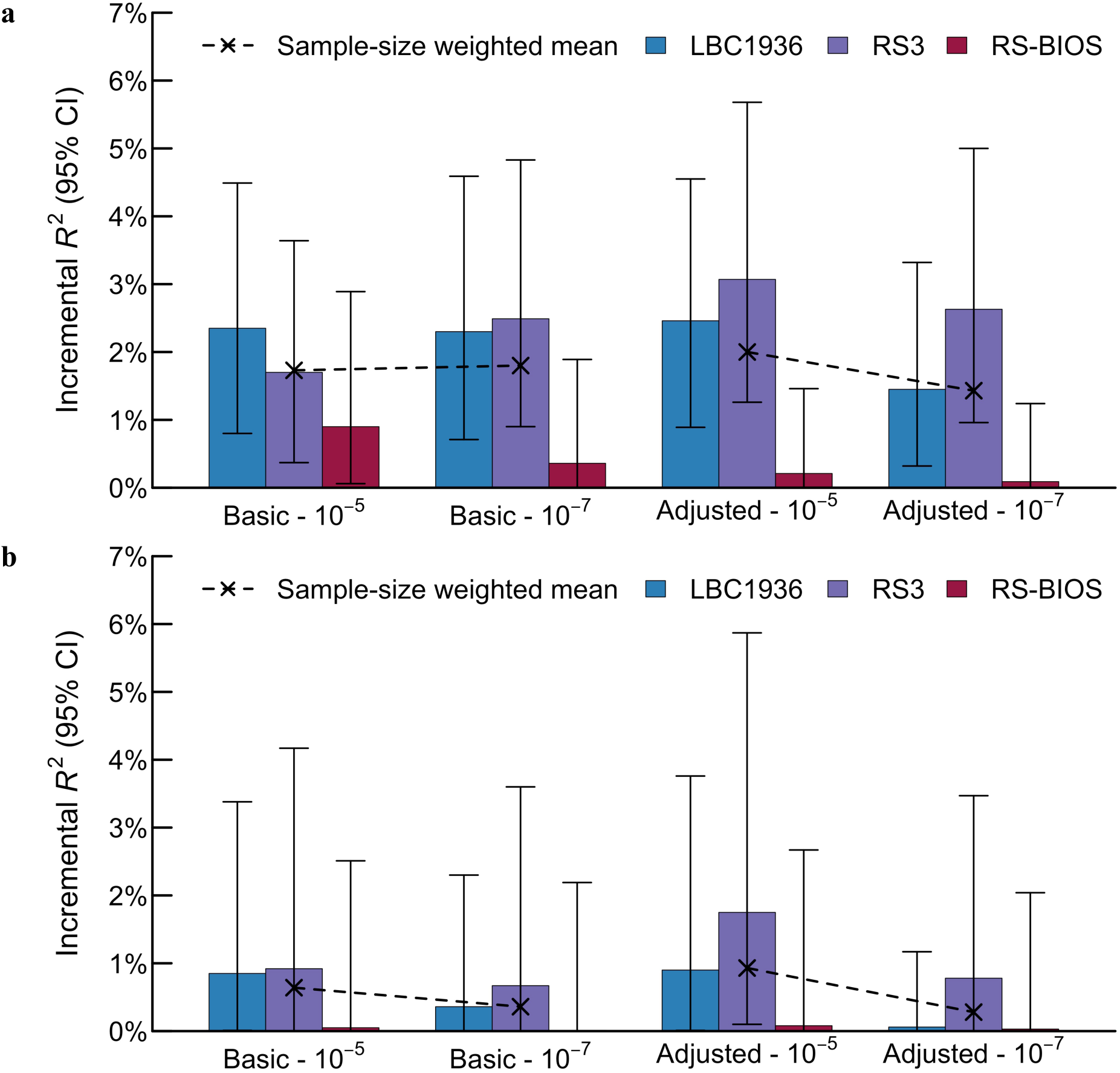
Methylation score prediction of educational attainment in independent holdout samples. Panel (a) displays the prediction in all individuals, and panel (b) displays the prediction restricted to never smokers. The incremental *R*^2^ of the methylation score was constructed with the coefficient estimates from meta-analysis of the basic model, as well as the adjusted model. The sample sizes of the LBC1936, the RS3, and the RS-BIOS cohorts are 918, 728, and 671 respectively. We performed sample size weighted meta-analysis across the cohorts for each of the 4 combinations of model estimates and *P*-value thresholds. The respective *P*-values are 4.42×10^−11^, 7.76×10^−11^, 2.02×10^−11^, and 3.28×10^−8^ respectively for the full sample and 0.0183, 0.0898, 0.0051, and 0.1818 respectively for the never-smokers. The full prediction results are presented in **Supplementary Table S1.9a and S1.9b.**

Relative to the prediction in the full sample, in the subsample of never-smokers the PGMSs (constructed with weights derived from the full EWAS sample) predict less of the variance in EA. As shown in **Figure 4** and **Supplementary Table S1.10b**, the incremental *R^2^*’s range from 0.3% to 0.9%. Two PGMSs constructed from probes reaching *P* < 1×10^−5^ in the EWAS are associated with EA at significance threshold *P* < 0.05, while the two PGMSs constructed only from the lead probes reaching *P* < 1×10^−7^ are not (P > 0.05). No interaction effect between the PGMS and the polygenic score is found in the never-smoker subsample.

Turning to the child sample in the ALSPAC ARIES cohort^44,47^, we examine whether a PGMS constructed from methylation assessed in cord blood samples at birth is predictive of four prospective measures of educational achievement test scores (Key Stage 1–4^45^), collected between ages 7 and 16 (**Supplementary Note 1.7.1**). The results are reported in **Supplementary Table S1.10c**. The highest incremental *R^2^* is 0.73% (*P* = 0.0094), and it is attained in the model predicting school performance at age 14-16 (i.e. the Key Stage 4 test scores). However, once maternal smoking status is added as a control variable, the predictive power of the PGMS is essentially zero (*R^2^* = 0.05%, *P* = 0.234). This suggests that the predictive power is driven by the confounding effects of maternal smoking. We draw two conclusions from these results from the child sample. First, they reinforce the concern that maternal smoking is a major confound for any probe associations with EA. Second, they suggest that any true methylation-EA associations are unlikely to be driven by a causal effect of methylation on EA.

#### Overlap between EWAS probes and published GWAS associations

To supplement our polygenic-score analyses of the overlap between epigenetic and genetic associations, we next investigated whether our lead probes are located in genomic regions that contain SNPs that have been identified in GWAS of EA or GWAS of smoking. Considering jointly the 141 approximately independent EWAS probes reaching *P* < 10^−4^, we did not find evidence of enrichment for either EA-linked SNPs (*P* = 0.206) or smoking-linked SNPs (*P* = 0.504) (see **Supplementary Note 2.4**). Considering the probes individually, one probe (cg17939805) was found to be in the same genomic region, with a genomic distance of 607 bp, as a SNP (rs9956387) associated with EA, whereas no probes were close to SNPs previously identified as linked to smoking.

#### Epigenetic clock associations with EA

Two cohorts, FINRISK and MCCS, did not contribute to the epigenetic clock analyses. Therefore, the sample sizes for these analyses are smaller than for the EWAS meta-analysis: 8,173 for the basic age acceleration model and 7,691 for the adjusted age acceleration model (the difference being due to the lack of covariates for some individuals). The results are presented in **Supplementary Table S1.11**. There is no evidence for an association between EA and Clocks 1, 2, or 3, but the association between EA and Clock 4 is strong (*P* = 3.51 ×10^−6^ and *P* = 4.51×10^−4^ in the basic and adjusted model, respectively). The point estimates are small, however: using Clock 4, each year of EA is associated with a 0.071-year (i.e., 26-day) reduction in age acceleration in the basic model and a 0.055-year (i.e., 20-day) reduction in the adjusted model. Overall then, higher educational attainment is associated with slightly younger biological age when compared with chronological age. We note that the epigenetic clock that is found to be associated with EA, Clock 4, has previously been found to be the most predictive epigenetic clock measure of mortality^46^.

## Discussion

This study provides one of the first large-scale investigations in humans of epigenetic changes linked to a biologically distal environmental factor. In our EWAS meta-analysis— one of the largest EWAS conducted to date—we found 9 CpG probes associated with EA. Each of these probes explains 0.3% to 0.7% of the variance in EA—effect sizes somewhat smaller than the largest EWAS effects that have been observed for BMI and many times smaller than those observed for alcohol consumption, smoking, and especially maternal smoking during pregnancy. When we restrict our analysis to the subsample of never-smokers, the effect sizes of 7 out of the 9 lead probes are substantially attenuated. Moreover, the other two lead probes have been found in previous work to be strongly associated with maternal smoking during pregnancy^12^. More generally, comparing our own results to those from previous EWAS highlights a variety of factors correlated with EA, including not only maternal smoking but also alcohol consumption and BMI, as potentially major confounds for the EA associations we detect. We also cannot rule out that other factors correlated with EA, such as exposure to second-hand smoke, could confound the EA associations.

Convincingly establishing a causal effect of EA would require analysing a sample with quasi-random variation in EA, such as a sample in which some individuals were educated after an increase in the number of years of compulsory schooling and other individuals were educated before the law change^48^. We are not aware of large EWAS samples with quasi-random variation at present, but we anticipate that such samples will become available as methylation becomes more widely measured.

Although the EWAS we report here is among the largest conducted to date, our sample size of 10,767 individuals is only large enough to identify nine probes associated with EA at the conventional epigenome-wide significance threshold. Subsequent EWAS conducted in larger samples that have sufficient statistical power to identify a much larger number of EA-associated probes will enable more extensive investigations of overlap with probes associated with other phenotypes than were possible from our results, as well as analyses of the biological functions of the probes. Besides limited statistical power, other limitations of our study, common to EWAS research designs, are that we study methylation cross-sectionally and not longitudinally and that we only investigate CpG methylation and not other types of epigenetic modifications.

## Conclusion

One plausible hypothesis is that environmental influences on the epigenome—even those due to everyday, social environmental factors—are pervasive and profound^3^. According to the logic of this view, a major life experience that occurs over many years, such as EA, should leave a powerful imprint on the epigenome. Motivated by this view and by the evidence of large EWAS effects in studies of lifestyle factors, when we embarked on this project we entertained the hypothesis that we might find large associations between EA and methylation. We also entertained the alternative hypothesis that EA, because it is so biologically distal, may exhibit much weaker associations with methylation.

While our results do not allow us to distinguish how much of the effects we find are due to true associations with EA and how much are due to confounds, they strongly suggest that the effect sizes we estimate are an upper bound on the effect sizes of any true methylation associations with EA. These upper-bound effect sizes are far smaller than associations with more biologically proximal environmental factors that have been studied. If our results can be generalized beyond EA to other biologically distal environmental factors, then they cast doubt on the hypothesis that such factors have large effects on the epigenome.

## ACKNOWLEDGEMENTS

This research was carried out under the auspices of the Social Science Genetic Association Consortium (SSGAC). The SSGAC seeks to facilitate studies that investigate the influence of genes on human behavior, well-being, and social-scientific outcomes using large genomewide association study meta-analyses. The SSGAC also provides opportunities for replication and promotes the collection of accurately measured, harmonized phenotypes across cohorts. The SSGAC operates as a working group within the CHARGE consortium. A full list of acknowledgments is provided in **Supplementary Note**. The authors declare no competing financial interests. Upon publication, results can be downloaded from the SSGAC website (http://thessgac.org/). Data for our analyses come from many studies and organizations, some of which are subject to a MTA, and are listed in the **Supplementary Note.** We thank Aysu Okbay for conducting the meta-analyses for the SNP polygenic scores. The data were accessed under Section 4 of the Data Sharing Agreement of the Social Science Genetic Association Consortium (SSGAC).

